# Inhibition of Chikungunya virus nsP2 protease *in vitro* by pantinin-1 isolated from scorpion venom

**DOI:** 10.1101/2025.06.29.662183

**Authors:** Mohammadamin Mastalipour, Mônika Aparecida Coronado, Jorge Enrique Hernández González, Dieter Willbold, Raphael Josef Eberle

## Abstract

Climate change has enhanced the spread of arboviruses such as Chikungunya virus (CHIKV). CHIKV is a re-emerging virus from the family *Togaviridae* that has spread globally, causing numerous outbreaks. The lack of antiviral therapy against CHIKV makes it a significant threat to public health. Cleavage of the viral polyprotein depends on the catalytic activity of nsP2, which is essential for viral replication. Due to this critical role, the nsP2 protease is a promising target for antiviral drug development. Animal venom-derived peptides have shown great potential against a variety of diseases, including infections, cancer, and neurodegenerative disorders. In this study, we evaluated the inhibitory effects and properties of pantinin-1, a peptide derived from the scorpion *Pandinus imperator* with broad antimicrobial activity, against CHIKV nsP2 protease. Pantinin-1 effectively inhibited CHIKV nsP2 protease, with a half-maximal inhibitory concentration (IC_50_) of 6.4 ± 2.04 µM and complete inhibition at 175 µM. Further analysis revealed that pantinin-1 acts as a competitive inhibitor with low micromolar affinity and showed no toxicity up to 20 µM in cell culture. Lastly, using molecular docking with subsequent molecular dynamics, the protein–peptide interaction was analyzed, and the key residues involved in the interaction with the protease were predicted. These findings highlight the potential inhibitory effect of pantinin-1 as a lead candidate targeting nsP2 protease.

## 1. Introduction

Arthropod-borne viruses (arboviruses) are a significant global health threat due to their ability to cause severe illness and widespread outbreaks (Grace et al., 2022). These viruses are primarily transmitted by arthropods such as mosquitoes and ticks and include human pathogens such as dengue virus (DENV), Zika virus (ZIKV), and chikungunya virus (CHIKV), all of which are transmitted by *Aedes* mosquito species (Delrieu et al., 2023). Among these viruses, CHIKV stands out due to its rapid geographic expansion, with cases reported in more than 100 countries across Africa, Asia, the Americas, and Europe (Li et al., 2021; Vairo et al., 2019; Zhang et al., 2025). CHIKV infection causes chikungunya fever, which typically presents in two clinical phases. The acute phase is characterized by high fever, headache, rash, and severe joint pain (Imad et al., 2021). In some patients, symptoms persist into a chronic phase, characterized by ongoing arthralgia, sleep disturbances, and psychological conditions such as depression (Paixao et al., 2018; Silveira-Freitas et al., 2024). Additionally, CHIKV has been associated with neurological complications, including encephalitis and peripheral neuropathies, particularly in neonates (Das et al., 2010).

CHIKV is a single-stranded, positive-sense RNA virus belonging to the *Togaviridae* family, genus *Alphavirus* (Wang et al., 2024). Its genome encodes four non-structural proteins (nsP1–nsP4) and five structural proteins including the capsid (C), envelope protein (E1,E2) and three accessory proteins (E3, transframe protein and 6k) (Ahola & Merits, 2016; Metz & Pijlman, 2016). Among these, non-structural protein 2 (nsP2) plays a key role in viral replication. It contains an RNA helicase domain with nucleoside triphosphatase (NTPase) and RNA triphosphatase activities, as well as a C-terminal protease and methyltransferase domain (Das et al., 2014). The nsP2 protease (nsP2^pro^) is a member of MEROPS clan CN, features a papain-like cysteine protease domain and is responsible for cleaving the viral polyprotein P1234 into individual functional units essential for replication (Ghoshal et al., 2024; Rausalu et al., 2016). Due to its essential enzymatic functions, nsP2^pro^ is a promising target for antiviral drug development. Several studies have explored potential inhibitors of nsP2^pro^ using *in silico, in vitro*, and biochemical approaches. These include peptide-based inhibitors such as P1 and natural products like hesperidin (Eberle et al., 2021; Mastalipour et al., 2025). However, none of these compounds have advanced beyond preclinical testing or received approval for human use. While two vaccines, VLA1553 (Schneider et al., 2023) and Vimkunya (Richardson et al., 2025), have recently been approved against CHIKV infection, vaccination is not effective for individuals who are already infected. This underscores the need for antiviral therapy.

In recent years, animal venoms from arthropods such as spiders and scorpions, as well as snakes and other venomous species have emerged as promising sources of bioactive compounds with pharmacological potential (Lyukmanova & Shenkarev, 2024). These venoms contain peptides and proteins with diverse mechanisms of action. They have been studied for applications including antimicrobial and antiviral therapies, as well as treatments for neurodegenerative diseases such as Alzheimer’s and Parkinson’s.(Alvarez-Fischer et al., 2013; Badari et al., 2020; Camargo et al., 2024; Zona Rubio et al., 2025). Besides the diversity and broad spectrum of venom-derived peptides, many of them exhibit advantages such as cell-penetrating properties, specificity, and resistance to enzymatic degradation (Lewis & Garcia, 2003; Rádis-Baptista, 2021). For example, a peptide from *Crotalus durissus terrificus* venom has demonstrated *in vitro* activity against amyloid-β_42_, a key pathological hallmark of Alzheimer’s disease (Camargo et al., 2024). In the context of infectious diseases, venom-derived peptides have shown broad-spectrum activity. Latarcins, derived from the spider *Lachesana tarabaevi*, have exhibited potent antibacterial and antiviral activity (Kozlov et al., 2006; Rothan et al., 2014). Likewise, the scorpion-derived peptide Mucroporin-M1 has shown virucidal activity against pathogens including SARS-CoV, measles virus, and influenza H5N1(Li et al., 2011). Additional venom-derived compounds with antimicrobial or antiviral activity are summarized in Table 1. In addition to these known peptides, pantinin-1 has gained attention due to its potent and broad-spectrum antimicrobial properties (Zeng et al., 2013). Pantinin-1, a 14-amino acid peptide derived from the scorpion *Pandinus imperator* (Crusca et al., 2018; Zeng et al., 2013). Pantinin-1 possesses a characteristic α-helical, amphipathic structure believed to underlie its membrane-disrupting mechanism and hemolytic activity (Zeng et al., 2013) It has demonstrated antimicrobial activity against Gram-positive bacteria such as *Staphylococcus aureus* (AB 94004) and MRSA (16472), Gram-negative strains such as *Escherichia coli* (DH5α) and *Klebsiella oxytoca* (AB 2010143), as well as antifungal activity against *Candida tropicalis* (AY 91009), with a minimum inhibitory concentration (MIC) of 16 µM (Zeng et al., 2013).

**Table 1.**
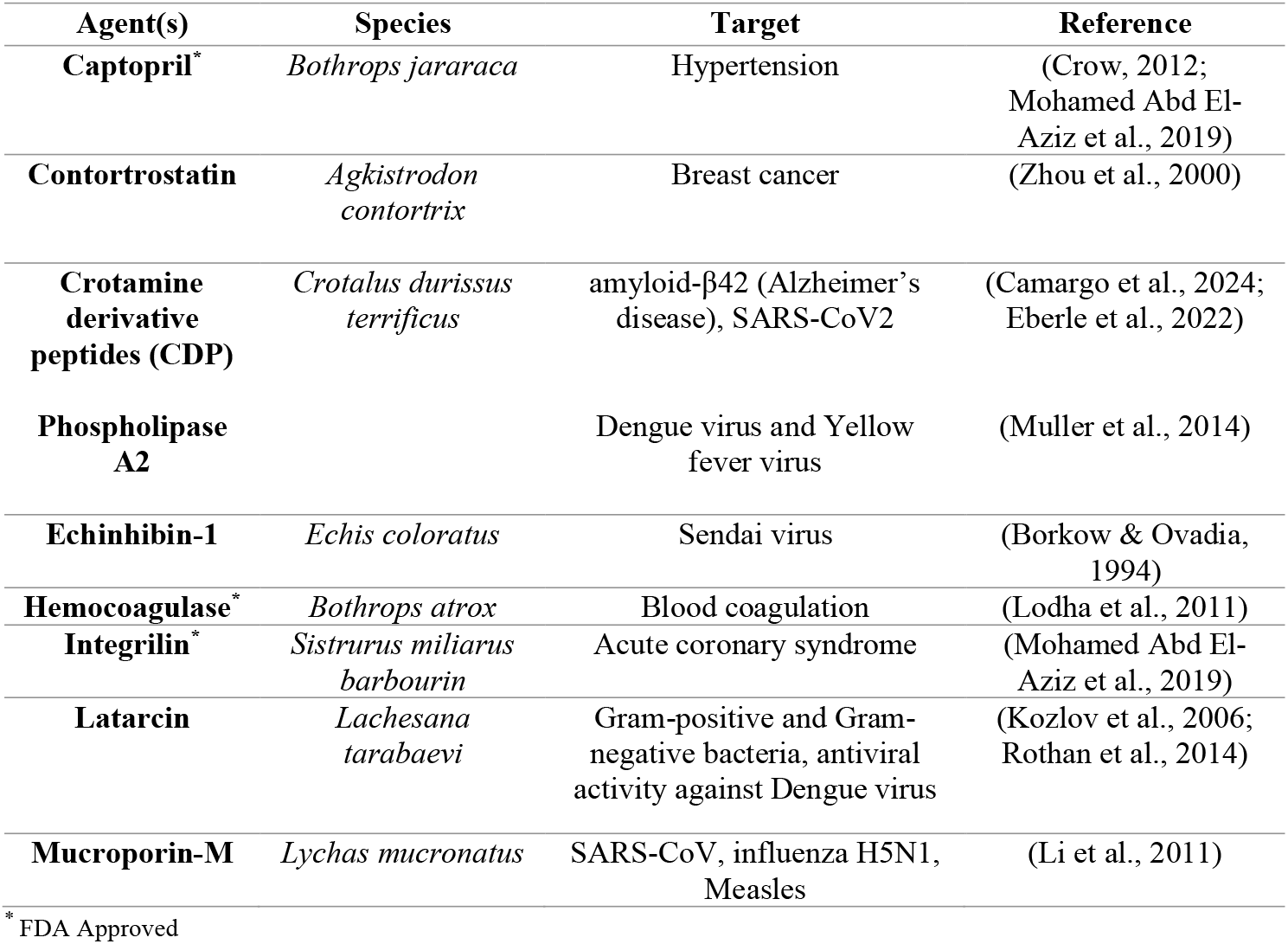
Representative proteins isolated from venom and venom-derived peptides with reported antimicrobial, antiviral, or therapeutic activity.

In this study, we investigated the inhibitory potential of pantinin-1 against the CHIKV nsP2 protease *in vitro*. Our findings revealed that pantinin-1 effectively inhibited nsP2^pro^ activity, with an IC_50_ of 6.4 ± 2.04 µM. Further analysis showed that pantinin-1 binds at the active site as a competitive inhibitor with an equilibrium dissociation constant (K_D_) of 9.29 μM. Cytotoxicity assessment using the MTT assay showed no detectable toxicity at concentrations up to 20 µM, while approximately 50% cell viability was observed at 40 µM. *In silico* studies predicted the most stable pose and interaction between the protease and pantinin-1, identifying key residues involved in binding. These results suggest that pantinin-1 has inhibitory effects within a concentration range that precedes the onset of cytotoxicity, supporting its potential as a lead compound for further development in CHIKV antiviral research.

## 2. Materials and methods

### 2.1 Protein expression and purification

Recombinant CHIKV nsP2 protease was expressed and isolated following a previously established and published protocol (Eberle et al., 2021; Mastalipour et al., 2025)

### 2.2 Peptide material

The peptide used in this study, pantinin-1, was supplied in solid, lyophilized form by GenScript Biotech (Rijswijk, Netherlands). It consists of 14 L-amino acids (GILGKLWEGFKSIV) and was chemically modified with an acetyl group at the N-terminus and an amide group at the C-terminus. Analytical documentation provided by the supplier confirmed that the peptide met a purity threshold exceeding 90%. Verification of peptide identity and purity was conducted by chromatographic and spectrometric methods. Reverse-phase HPLC was performed using an Inertsil ODS-0310 analytical column (dimensions: 4.6 mm × 250 mm), and molecular weight confirmation was achieved via electrospray ionization mass spectrometry (ESI-MS). The full analytical documentation supporting peptide purity and identity is shown in the Supplementary Information (Fig. S1).

### 2.3 Enzymatic inhibition assay of CHIKV nsP2^pro^

The inhibitory effect of pantinin-1 on CHIKV nsP2^pro^ was assessed using a fluorescence-based enzymatic assay. This assay employed a synthetic peptide substrate labeled with DABCYL and EDANS fluorophores (DABCYL-Arg-Ala-Gly-Gly-↓Tyr-Ile-Phe-Ser-EDANS; BACHEM, Bubendorf, Switzerland), which reflects the native cleavage sequence of the CHIKV polyprotein (Eberle et al., 2021b; Hu et al., 2016). The assay was conducted in a 96-well microplate with a total volume of 100 µL per well. Each reaction mixture contained 20 mM Bis-Tris propane buffer (pH 7.5), 10 µM nsP2^pro^, and 9 µM fluorogenic substrate. Pantinin-1 was prepared as a 10 mM stock solution in DMSO and diluted into the reaction mixture to achieve final concentrations ranging from 0 to 175 µM. Fluorescence emission, indicative of substrate cleavage, was monitored at 37 °C using a CLARIOstar plate reader (BMG Labtech, Ortenberg, Germany) with excitation and emission wavelengths set to 340 nm and 490 nm, respectively. Readings were taken at 30-second intervals for 30 minutes. Enzymatic activity in the presence of pantinin-1 was expressed as a percentage of the activity observed in untreated control samples, calculated using the equation (E1) (Eberle et al., 2021).

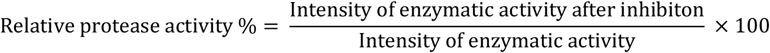

To determine the IC_50_ value, non-linear regression analysis was performed using GraphPad Prism (version 5). All experiments were performed as triplicates, and the results are presented as mean ± standard deviation (SD).

### 2.4 Inhibition mechanism analysis

To characterize how pantinin-1 interacts with the catalytic function of CHIKV nsP2^pro^, an inhibition assay was performed using the same FRET-based platform described previously. The enzyme concentration was kept constant at 10 µM, while both substrate and inhibitor levels were systematically varied. All reactions were carried out in 20 mM Bis-Tris-Propane buffer, pH 7.5. The resulting data, which include combinations of multiple substrate and inhibitor concentrations (Table 2), were used to evaluate the inhibition pattern using Lineweaver–Burk plot.

**Table 2.** Substrate and peptide concentrations used in the inhibition-mode assay.

| Substrate conc. [μM] | 0 | 1 | 2.5 | 5 | 7.5 | 10 |
| --- | --- | --- | --- | --- | --- | --- |
| Inhibitor conc. [μM] | 0 | 0 | 0 | 0 | 0 | 0 |
|  | 1 | 1 | 1 | 1 | 1 | 1 |
|  | 2.5 | 2.5 | 2.5 | 2.5 | 2.5 | 2.5 |
|  | 5 | 5 | 5 | 5 | 5 | 5 |

### 2.5 Biolayer interferometry (BLI)

Biolayer interferometry (BLI) was used to determine the equilibrium dissociation constant (K_D_) of pantinin-1. Experiments were performed using an Octet RED96 system (Sartorius, Göttingen, Germany) with Octet AR2G biosensors (Sartorius, Göttingen, Germany). The biosensors were incubated in running buffer (20 mM Bis-Tris propane, 2% DMSO, pH 7.5) for 15 min before starting the experiment. The experiments were initiated with an equilibration step with running buffer for 600 s. Next, the sensor surface was activated with activation buffer containing 100 mM NHS and 50 mM MES, pH 5.2 (Xantec, Düsseldorf, Germany), for 420 s. The nsP2^pro^ (10 µg/mL) was then immobilized onto the activated biosensor surface for 420 s in 20 mM sodium acetate, pH 5.0. The reaction was then quenched using 1 M ethanolamine-HCl, pH 8.5 (Xantec, Düsseldorf, Germany), for 300 s. Afterwards, sensors were re-equilibrated in running buffer for an additional 600 s. A six-step 1:3 serial dilution of pantinin-1 was prepared, ranging from 20 µM to 0 µM. Each sensor was exposed to pantinin-1 during an association step for 180 s, followed by a 600 s dissociation phase. After each cycle, sensors were regenerated with 20 mM glycine (pH 2.0) and equilibrated again in running buffer for 600 s. All experiments were performed in triplicate. To investigate the binding of pantinin-1 on sensors, a control experiment was performed same as described above without immobilizing the protease, with the highest concentrations of pantinin-1 at 20 µM and 6.66 µM. Data were analyzed using ForteBio Data Analysis Software 8.0 (Sartorius, Göttingen, Germany). Binding responses from three independent experiments were combined to construct a Scatchard plot, from which the dissociation constant (K_D_) was determined (Kessler et al., 2008).

### 2.6 Cell viability assay

The cytotoxic potential of pantinin-1 was evaluated in Vero cells (African green monkey kidney epithelial cell line, *Chlorocebus aethiops*). Cells were cultured in Dulbecco’s Modified Eagle Medium (DMEM) supplemented with 10% fetal calf serum (FCS) and 1% non-essential amino acids, and maintained at 37 °C in a humidified incubator with 5% CO_2_. For the assay, cells were seeded in 96-well plates and treated with pantinin-1 at final concentrations ranging from 0 to 100 µM. The peptide was dissolved in DMSO with final concentration of 10 mM and subsequently diluted in complete culture medium. To establish a control for maximal cytotoxicity, 0.1% Triton X-100 was used as a positive control. After overnight incubation with the test compound, cell viability was assessed using the MTT assay (Roche Diagnostics GmbH, Mannheim, Germany). 10 µl of MTT reagent were added to each well, followed by a 4 h incubation to allow for formazan formation by metabolically active cells. Crystals were dissolved by the addition of 100 µL of solubilization solution, and the plates were incubated overnight to ensure complete dissolution. Absorbance was recorded at 570 nm and 660 nm using a CLARIOstar plate reader (BMG Labtech, Ortenberg, Germany). Cell viability was calculated using the following equation:

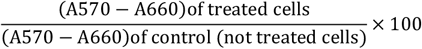

### 2.7 Protein-peptide docking

An MD-derived nsP2^pro^ conformation with an open active site, reported in a previous work (Mastalipour et al., 2025), was employed here to dock the pantinin-1 peptide. The three-dimensional structure of pantinin-1 used for docking was predicted with AlphaFold3 (Abramson et al., 2024). Three docking programs, Galaxy TongDock (Park et al., 2019), ClusPro (Kozakov et al., 2017) and HDock (Yan et al., 2017), were used for this purpose with the default parameters established in their respective web servers. The best poses plus three poses exhibiting high conformational variation (i.e., large pair-wise peptide RMSD values) among the top-5 solutions determined with each docking program were selected for subsequent analyses to predict the most likely binding mode of pantinin-1 to nsP2^pro^, as described below.

### 2.8 MD simulations

The selected nsP2^pro^-pantinin-1 complexes were prepared for MD simulations with tleap of Amber22 (Case et al., 2022) as described in Mastalipour et al. (Mastalipour et al., 2025). Briefly, the peptide was capped at the N- and C-termini with ACE and NME moieties. The parameters for both the peptide and the protein were drawn from the ff19SB force field (Tian et al., 2020). Octahedral boxes were defined to simulate each system, with edges placed at least 10 Å away from the solute surface. The simulation boxes were then filled with OPC water molecules (Izadi et al., 2014) and 16 Cl-counterions were added to neutralize the solute net charge. Each system underwent energy minimization, followed by NVT heating and equilibration at the NPT ensemble to ensure a final temperature and pressure of 300 K and 1 bar, respectively. Then, the stepwise decrease of the harmonic restraints applied to the solute’s heavy atoms during the equilibration phase was conducted during four NPT simulations. Finally, 1 μs productive runs were run for every complex. All the MD simulations were performed with pmemd.cuda of Amber22 (Case et al., 2022). More details on the MD simulation setup can be found elsewhere (Mastalipour et al., 2025).

### 2.9 MM-GBSA free energy calculations

Molecular Mechanics Generalized-Born Surface Area (MM-GBSA) calculations were performed for the three top-ranked poses of peptide P1 in complex with CHIKV nsP2^pro^ using the MMPBSA.py module of Amber22 (Case et al., 2022; Miller et al., 2012), following the protocol detailed in (Mastalipour et al., 2025). The single-trajectory approach was employed, extracting 542 frames from the final 0.5 μs of each MD trajectory. The GB-neck2 model (igb = 8) (Nguyen et al., 2013) was used to estimate polar solvation energies, with internal and external dielectric constants set to 1 and 80, respectively, and a salt concentration of 0.1 M. Per-residue free energy decomposition was also performed under the same conditions to identify key binding interface residues (Mastalipour et al., 2025).

### 2.10 Trajectory analyses

Trajectory analyses were conducted using the cpptraj module from Amber22 (Case et al., 20022). Root-mean-square deviation (RMSD) values were obtained via the rms command. Clustering analysis employed the cluster command with a hierarchical agglomerative algorithm, using RMSD of peptide heavy atoms as the distance metric. Typically, five clusters were generated, and the most populated cluster was selected for structural representation. Hydrogen bond analysis was performed with the hbond command, applying geometric criteria of a donor–hydrogen–acceptor angle greater than 120° and a hydrogen–acceptor distance of 3.2 Å or less to define intramolecular hydrogen bonds.

### 2.11 Statistical Analysis

All statistical analyses were conducted using GraphPad Prism version 5.0. Differences between treatment groups and the control were evaluated using one-way analysis of variance (ANOVA) followed by Tukey’s post hoc test for multiple comparisons. Statistical significance was denoted as follows: p < 0.05 (*), p < 0.01 (**), and p < 0.001 (***).

## 3. Result

### 3.1. Inhibitory effect of pantinin-1 on CHIKV nsP2^pro^ activity

CHIKV nsP2^pro^ was expressed and purified as previously described (Eberle et al., 2021; Mastalipour et al., 2025). The pure protease was used to test the inhibitory potential of α-helical, amphipathic pantinin-1 (Fig. 1A) against the proteolytic activity of the CHIKV nsP2^pro^, which was assessed using a FRET-based enzymatic assay. Enzymatic activity was determined by measuring fluorescence emission over a 30-min period at excitation and emission wavelengths of 340 and 490 nm, respectively. As shown in Fig. 1B, pantinin-1 exhibited a clear, concentration-dependent inhibitory effect on nsP2^pro.^ A reduction in enzymatic activity was observed at low micromolar concentrations, with approximately 50% inhibition at around 5 µM. This suggests that pantinin-1 is capable of efficiently interfering with nsP2^pro^ activity at relatively low concentrations. The inhibitory effect became progressively stronger with increasing peptide concentration, culminating in a complete loss of enzymatic activity at 175 µM, the highest concentration tested. To quantify the potency of pantinin-1, a dose–response curve was generated, and the IC_50_ value was calculated using non-linear regression analysis in GraphPad. As shown in Fig. 1C, the IC_50_ was determined to be 6.4 ± 2.04 µM, based on three independent replicates

**Fig. 1.**
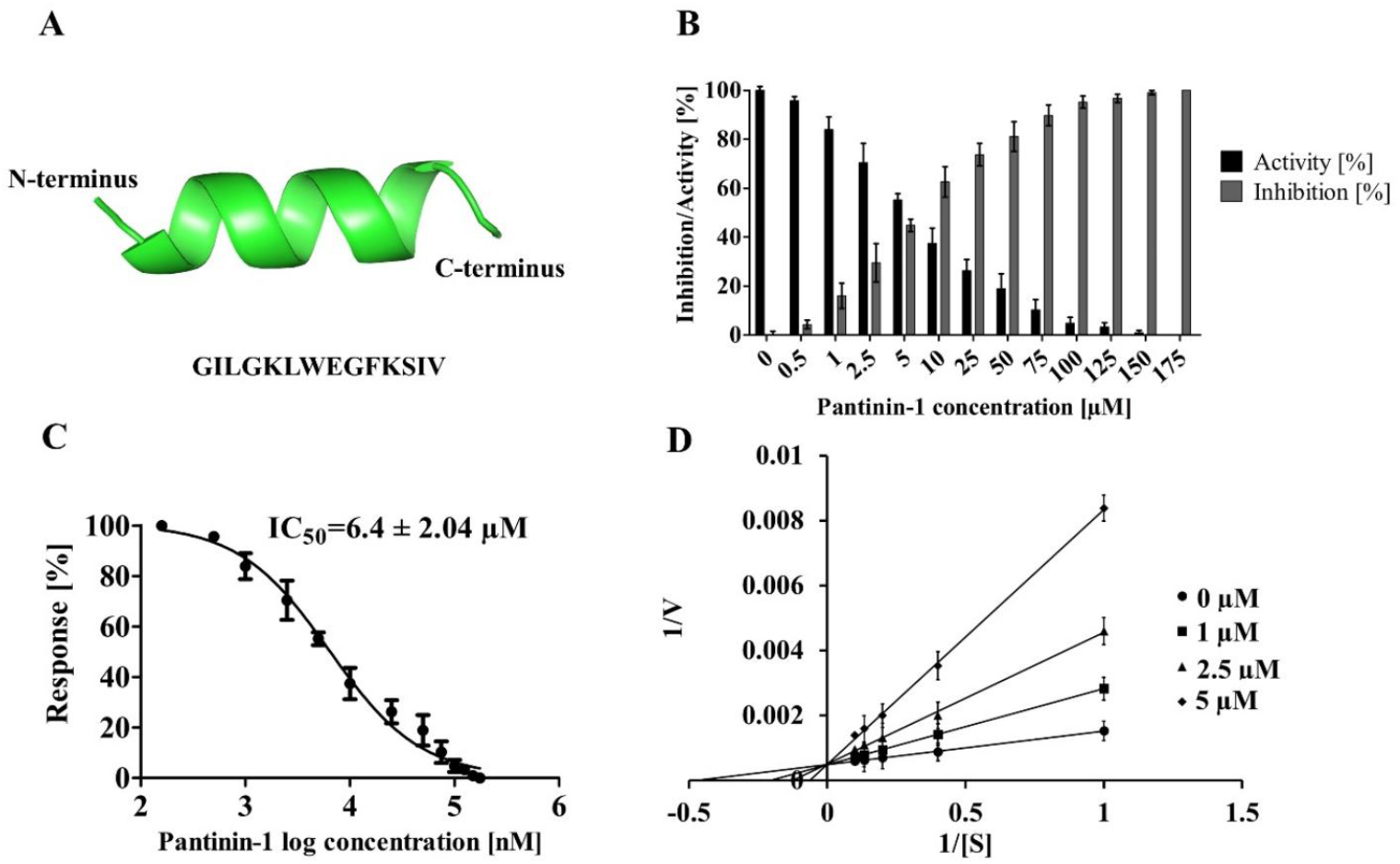
Pantinin-1 structure prediction and its inhibitory activity and binding analysis of pantinin-1 toward CHIKV nsP2^pro^. **(A)** The predicted structure of pantinin-1 generated by AlphaFold3 **(B)** Concentration-dependent inhibition of CHIKV nsP2^pro^ activity by pantinin-1, measured using a fluorescence-based enzymatic assay. Fluorescence was recorded over 30 min at 340/490 nm and 37 °C. Enzymatic activity was expressed as a percentage relative to the untreated control. **(C)** Dose–response curve of pantinin-1 inhibition, generated by non-linear regression analysis using GraphPad Prism. The calculated IC_50_ value was 6.4 ± 2.04 µM (mean ± SD; n = 3 technical replicates). **(D)** Kinetic evaluation of the inhibition mechanism of pantinin-1 using a Lineweaver–Burk plot. The intersection of the lines on the y-axis is characteristic of a competitive inhibition pattern, indicating that pantinin-1 competes with the substrate for binding to the enzyme’s active site. This results in an apparent increase in the Michaelis constant (K_m_).

### 3.2 Mode of inhibition of pantinin-1 toward CHIKV nsP2^pro^

In Section 3.1, we demonstrated the inhibitory effect of pantinin-1 on the enzymatic activity of CHIKV nsP2^pro^. To further characterize this interaction, it was essential to investigate the mechanism and mode of inhibition of pantinin-1. To this end, a FRET-based enzymatic assay was conducted using varying concentrations of both substrate and pantinin-1. The resulting data were analyzed using a Lineweaver– Burk plot, which allows assessment of how the inhibitor affects substrate binding. As shown in Fig. 1D, increasing the concentration of pantinin-1 at multiple fixed substrate concentrations led to a progressive increase in the apparent K_m_ constant. This pattern indicates that pantinin-1 decreasing the substrate’s binding affinity to nsP2^pro^.

### 3.3 Equilibrium dissociation constant measurement by biolayer interferometry (BLI)

To determine the equilibrium dissociation constant (K_D_) for the interaction between pantinin-1 and CHIKV nsP2^pro^, biolayer interferometry (BLI) was used. The experiment was performed using an Octet RED96 system (Sartorius, Göttingen, Germany) equipped with AR2G biosensors. Pantinin-1 was tested at six different concentrations, ranging from 20 µM to 0 µM. A sensor exposed only to the running buffer during the assay as the reference (0µM). All experiments were conducted at room temperature. The assay consisted of an association phase of 180 s followed by a dissociation phase of 600 s. A representative sensorgram is shown in Fig. S2A. All measurements were conducted in triplicate. From the BLI binding response data obtained across six concentrations of pantinin-1, a Scatchard plot was constructed using values extracted from ForteBio Data Analysis Software. This analysis yielded a dissociation constant (K_D_) of 9.29 μM (Fig. S2B).To ensure that the observed binding was specifically between the protease and pantinin-1, a control experiment was performed in the absence of the protease using 20 µM and 6.66 µM pantinin-1. As shown in Fig. S3, no binding was observed between the sensor and the peptide, indicating that the interaction occurs specifically between the protease and pantinin-1.

### 3.4 Cytotoxicity assessment of pantinin-1 in Vero cells

To investigate the cytotoxic potential of pantinin-1, Vero cells were exposed to a concentration gradient (0–100 µM) of the peptide, which was initially dissolved in DMSO and subsequently diluted in cell culture medium. Cell viability was determined using the MTT assay. As illustrated in Fig. 2, pantinin-1 exhibited no detectable cytotoxicity at concentrations up to 20 µM, with viability levels comparable to untreated controls. A notable decrease in viability occurred at 40 µM, where approximately half of the cell population remained viable. Exposure to higher concentrations (60–100 µM) resulted in a progressive and substantial loss of cell viability, with near-complete cytotoxicity observed at 80 µM and above.

**Fig. 2.**
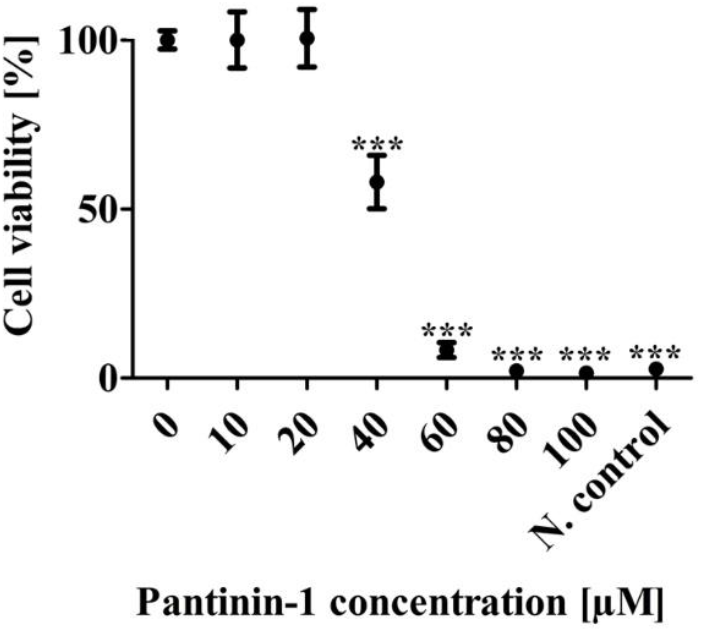
Cytotoxicity of pantinin-1 in Vero cells assessed by MTT assay. Pantinin-1 was dissolved in DMSO and diluted in culture medium to final concentrations ranging from 0 to 100 µM. Vero cells were exposed to the peptide, and cell viability was assessed using the MTT assay. No cytotoxic effects were observed at concentrations up to 20 µM. At 40 µM, viability decreased to approximately 50%, and at higher concentrations, a further decline was observed, with most cells non-viable at 100 µM. 0.1% Triton X-100 was used as a negative control. Data are presented as mean ± SD from three independent experiments. Statistical significance was determined by one-way ANOVA, with p < 0.001 considered significant. Asterisks indicate statistically significant differences compared to untreated controls.

### 3.5 Prediction of the nsP2^pro^-pantinin-1 complex structure

Protein-peptide docking was performed to predict the binding mode of pantinin-1 to nsP2^pro^. The top-5 poses determined with the three docking algorithms chosen in this work are shown in Fig. 3. Most of the solutions point at the CHIKV nsP2^pro^ active site as the region preferentially targeted by pantinin-1. However, ClusPro and HDock also predicted alternative binding modes, with pantinin-1 interacting at the interdomain hinge region located opposite the active site (Figs. 3B-C).

**Fig. 3.**
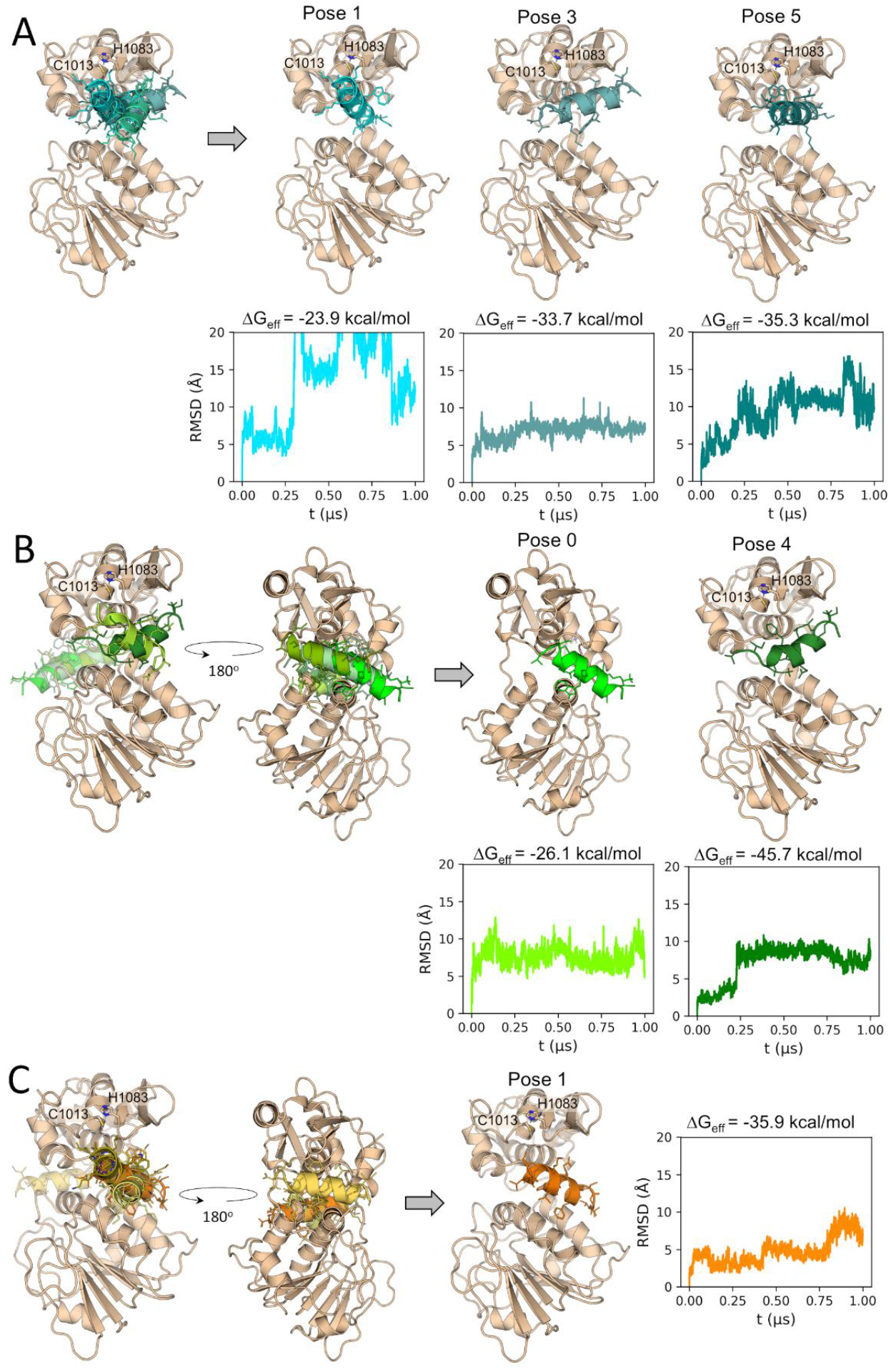
Prediction of the CHIKV nsP2^pro^-pantinin-1 complex through docking, MD simulations and free energy calculations. Superimposed five top-scoring docking poses of pantinin-1 predicted, determined with **(A)** Galaxy TongDock, **(B)** ClusPro and **(C)** HDock. Docking poses selected for MD simulations are depicted after the gray arrows. The RMSD time profiles of pantinin-1 from the simulated complexes are shown using the same colors as the corresponding peptide poses. The effective binding free energies (ΔG_eff_) calculated from the MD simulations are shown above each RMSD graph. RMSD values were calculated for the peptide backbone atoms and with respect to the peptide’s conformation in the docking pose. Prior to peptide RMSD calculations, all trajectory frames were superimposed on the nsP2^pro^ backbone in the initial conformation. The catalytic residues C1013 and H1083 are represented as sticks and labeled in the structural representation of the different CHIKV nsP2^pro^-pantinin-1 docking poses.

As described in Materials and Methods, the best pose predicted by each docking algorithm, plus three poses selected among the top-5 poses generated by each algorithm, were chosen for subsequent analyses to determine their stability. The peptide RMSD values calculated during the MD simulations of the selected docking poses indicate that at least in three systems, i.e., Galaxy TongDock pose3, and ClusPro poses 0 and 4, that the peptide reached stable conformations, characterized by RMSD plateau regions (Fig. 3). Moreover, the calculated ΔG_eff_ values for all the simulated systems predict the conformation adopted during the second half of the MD simulation initiated from the ClusPro pose 4 as the most stable one.

The representative structure calculated for the trajectory of the most stable pose in complex with CHIKV nsP2^pro^ (ClusPro pose 4, Fig. 3B), is depicted in Fig. 4A. As can be observed, the final pantinin-1 binding mode diverges significantly from the docking pose, as also deduced by the peptide RMSD profile (Fig. 3B and 4A). At the new position, the peptide occupies an active site pocket at the interdomain region, while leaving the catalytic residues exposed. The free energy decomposition suggests a dominant contribution of hydrophobic interactions, as deduced from the nature of the most important residues at the interface (Fig. 4B). Residues L3, W7, F10, I13 and V14 of pantinin-1 are predicted as key residues for nsP2^pro^ binding. Their mutation to Ala can serve as a strategy to experimentally validate the conclusions derived from the *in silico* analyses presented here.

**Fig. 4.**
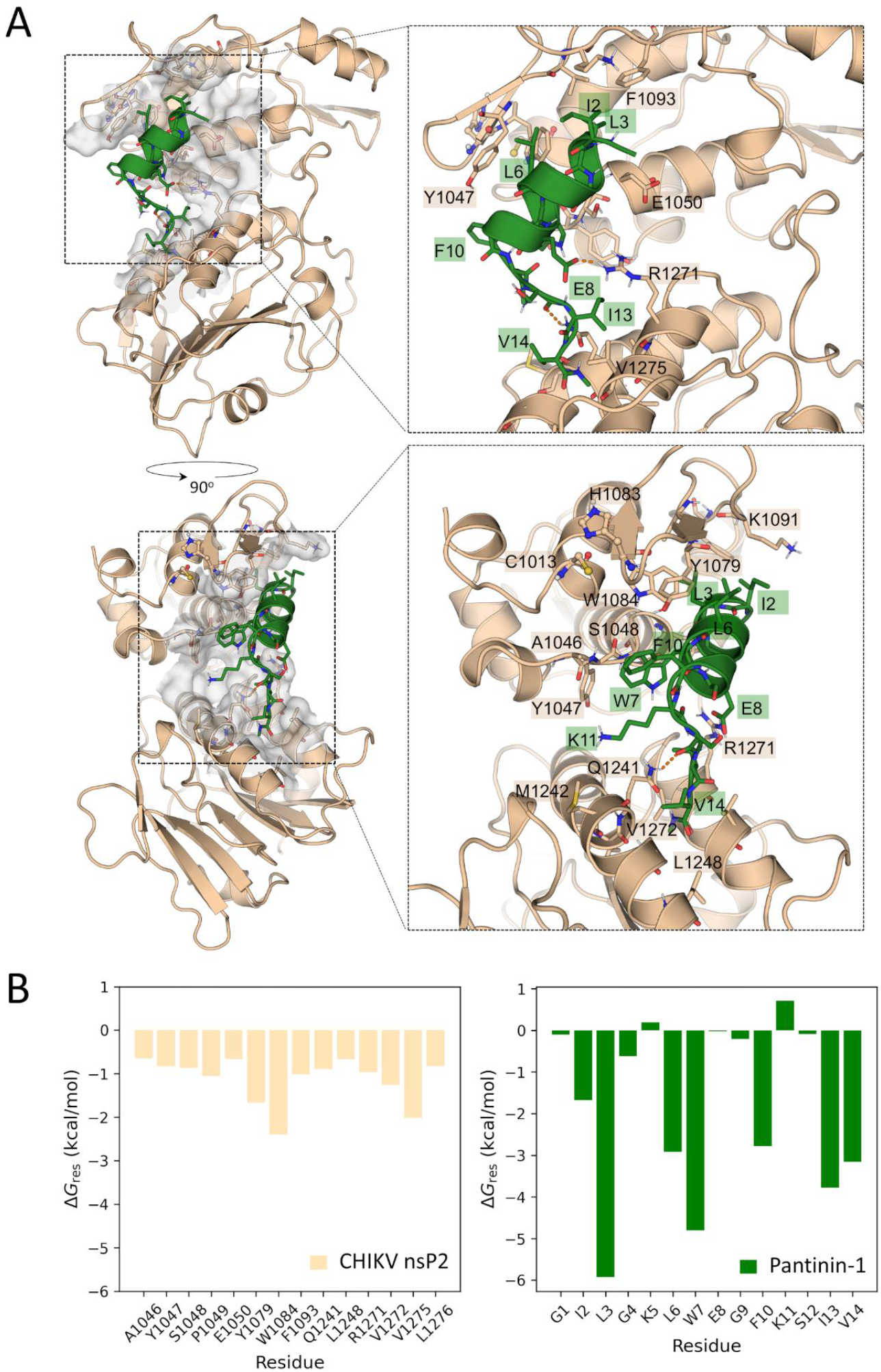
Representative structure of the predicted CHIK nsP2^pro^:pantinin-1 complex and the energy contribution of residues at the interface. **(A)** Two views of the CHIKV nsP2^pro^:pantinin-1 complex. Interface residues are shown in sticks and are labeled. The active site residues C1013 and H1083 are represented as sticks and spheres. H-bonds are indicated as orange dashed lines. **(B)** Per-residue free energy contributions of the nsP2^pro^ interface residues (<0.6 kcal/mol) and of all pantinin-1 residues.

## 4. Discussion

Chikungunya virus (CHIKV), along with Dengue, Yellow Fever, and Zika viruses, is among the major emerging arboviruses transmitted by *Aedes* species mosquitoes and is responsible for outbreaks worldwide (Barreto-Vieira et al., 2021; Grace et al., 2022). CHIKV belongs to the genus *Alphavirus* (Delrieu et al., 2023), and its replication depends on the multifunctional non-structural protein 2 (nsP2). The C-terminal region of nsP2 contains a protease domain (nsP2^pro^), which plays a crucial role in processing the viral polyprotein, an essential step in the viral life cycle (Rausalu et al., 2016). Over the years, various strategies have been explored to inhibit CHIKV nsP2^pro^ as a potential antiviral target. Techniques such as phage display have been used to identify inhibitory compounds (Mastalipour et al., 2025). Although numerous candidates have been reported, none have advanced beyond the laboratory stage (Table 3).

**Table 3.** Summary of characterized CHIKV nsP2^pro^ inhibitors from prior research.

| Compound | Type | $\text{IC}_{50}$ | $K_D$ ($\mu\text{M}$) | Inhibition Mode | Reference |
| --- | --- | --- | --- | --- | --- |
| **Peptide P1** | Peptide | $4.6 \pm 1.9 \mu\text{M}$ | $1.39 \pm 0.61$ | Competitive | (Mastalipour et al., 2025) |
| **RA-0002034** | Small molecule | $58 \pm 17 \text{ nM}$ | N/A | Covalent (Cys binding) | (Merten et al., 2024) |
| **Withaferin A (WFA)** | Natural compound | $0.51 \mu\text{M}$<br>(In BHK-21 cells) | 64 | N/A | (Sharma et al., 2024) |
| **Hesperetin (HST)** | Natural flavonoid | $2.5 \pm 0.4 \mu\text{M}$ | $31.6 \pm 2.5$ | Noncompetitive | (Eberle et al., 2021) |
| **Hesperidin (HSD)** | Natural flavonoid | $7.1 \pm 1.1 \mu\text{M}$ | $40.7 \pm 2.0$ | Noncompetitive | (Eberle et al., 2021) |
| **Pep-I** | peptidomimetic | 34 $\mu\text{M}$ | N/A | Noncompetitive | (Singh et al., 2018) |
| **Pep-II** | peptidomimetic | 42 $\mu\text{M}$ | N/A | Competitive | (Singh et al., 2018) |
| **MBZM-N-IBT** | Small Molecule | 31.96 | $6.14 \pm 0.58$ | Competitive | (De et al., 2022) |

Venom-derived peptides are new approach to develop a new therapeutic agent. These peptides have shown potential in treating a wide range of diseases, including bacterial and viral infections, cancer, and neurodegenerative disorders. For example, Phospholipase A2, isolated from the venom of *Crotalus durissus terrificus*, has shown antiviral activity against Dengue and Yellow Fever viruses (Muller et al., 2014). Ziconotide, a peptide from the marine snail *Conus magus*, is approved for chronic pain management (Deer et al., 2019; Rauck et al., 2009). Mastoparan and its synthetic analogs, derived from wasp venom, exhibit antiviral activity against Human *alphaherpesvirus 1* (HSV-1) (Vilas Boas et al., 2024). Similarly, Dermaseptins from hylid frogs have demonstrated broad-spectrum antimicrobial and anti-tumor properties (Bartels et al., 2019). Another venom-derived candidate is pantinin-1, a 14-amino-acid peptide isolated from the scorpion *Pandinus imperator*, known for its antibacterial activity (Zeng et al., 2013).

In this study, we evaluated pantinin-1 as a potential inhibitor of CHIKV nsP2^pro^. A FRET-based assay was conducted using pantinin-1 at concentrations ranging from 0 to 175 µM to determine inhibition effect of the pantinin-1. As shown in Fig. 1B, increasing concentrations of pantinin-1 progressively inhibited protease activity, with complete inhibition observed at 175 µM. Dose–response analysis (Fig. 1C) revealed an IC_50_ of 6.4 ± 2.04 µM. This potency is comparable to that of other known inhibitors (Table 3) such as Peptide P1 (4.6 µM) (Mastalipour et al., 2025), Hesperetin (2.5 µM) (Eberle et al., 2021), and Hesperidin (7.1 µM) (Eberle et al., 2021), although it is higher than that of RA-0002034, which exhibited a much lower IC_50_ of 58 ± 17 nM (Merten et al., 2024).

To understand the mechanism of inhibition, we performed Lineweaver–Burk analysis using varying concentrations of both substrate and pantinin-1. The resulting plot (Fig. 1D) showed all lines intersecting at the y-axis, with increased K_m_ and unchanged V_max_ indicating that pantinin-1 acts as a competitive inhibitor (Kenakin, 2017). This mode of inhibition is consistent with what we previously observed for peptide P1 (Mastalipour et al., 2025). Notably, both pantinin-1 and peptide P1 share a high content of α-helical secondary structure, which we have previously suggested may play a key role in peptide binding to the nsP2^pro^ active site. Next, we evaluated the binding affinity of pantinin-1 to CHIKV nsP2^pro^ using biolayer interferometry (BLI). Binding responses were measured across a concentration range of 0–20 µM, and values extracted from three independent experiments were used to construct a Scatchard plot. As shown in Fig. S2B, this analysis yielded a dissociation constant (K_D_) of 9.29 µM. While the binding affinity of pantinin-1) is slightly weaker than that of peptide P1 (1.39 µM) (Mastalipour et al., 2025), it is significantly stronger than that of Hesperetin (31.6 ± 2.5µM) and Hesperidin (40.7 ± 2.0 µM), as reported by Eberle et al. (Eberle et al., 2021). This difference does not necessarily indicate that pantinin-1 has superior binding, as various methods were used to determine the K_D_ of the inhibitors. For example, in the case of peptide P1, microscale thermophoresis was used, which underscores that this difference in affinity may lie in the different methods used.

We also assessed the cytotoxicity of pantinin-1 on Vero cells across a concentration range of 0–100 µM. Fig. 2 shows that pantinin-1 had no detectable cytotoxic effect up to 20 µM. At 40 µM, cell viability decreased to around 50%, and at 80-100 µM, most cells were non-viable. As mentioned before, pantinin-1 is an α-helical, amphipathic peptide (Zeng et al., 2013). It exhibits both antimicrobial and hemolytic activity by interacting with and disrupting cell membranes (Zeng et al., 2013). This property explains the toxicity observed in the MTT assay, where cell death occurred at higher concentrations of pantinin-1. In addition, Zeng et al. reported that the hemolytic activity of pantinin-1 begins at concentrations above 32 µM, which is consistent with the cytotoxic effects observed in this study (Zeng et al., 2013). Although cytotoxicity occurs only at concentrations above 20 µM, this still represents more than three times of the IC_50_.

During the molecular docking studies with three different programs (ClusPro (Kozakov et al., 2017) HDock (Yan et al., 2017) and Galaxy TongDock (Park et al., 2019)), we were able to predict likely conformations for the protease/pantinin-1 complexes. Most of these complexes showed binding at the active site of the protease (Fig. 3A). However, two of the programs exhibited conformations among the top-5 poses in which pantinin-1 binds to a region opposite to the active site (Fig. 3B–C). Further analysis, consisting in MD simulations followed by MM-GBSA free energy calculations, led to the identification of the most stable binding mode of pantinin-1 to CHIKV nsP2^pro^ (Pose 4; Fig. 3B). In the identified conformation, pantinin-1 interacts with active site residues located in both domains of the protease (Fig. 4A). The proposed binding mode is consistent with the competitive inhibition mechanism determined for this peptide.

Free energy decomposition analysis demonstrated that hydrophobic interactions are the dominant force between pantinin-1and CHIKV nsP2^pro^. The lowest per-residue free energies (ΔG_res_) correspond to L3, W7, F10, I13, and V14, indicating their important role in the interaction. However, this result needs to be confirmed, for example, by alanine substitution. Moreover, as shown in Fig. 4B, residues Y1079, Y1047, and W1084 are involved in the interaction with Pantinin-1. These amino acids are part of the substrate-binding site subunits S2, S3, and S4, suggesting possible binding at the substrate-binding site (Narwal et al., 2018). Overall, we demonstrated that pantinin-1, in addition to its antimicrobial activity, exhibits an inhibitory effect against CHIKV nsP2^pro^ *in vitro*, with binding affinity and an IC_50_ in the low micromolar range. This makes pantinin-1 a promising candidate for antiviral therapy development

## 5. Conclusion and outlook

In this study, it was demonstrated that pantinin-1, a peptide derived from scorpion, has the potential to inhibit CHIKV nsP2^pro^ with an IC_50_ of 6.4 ± 2.04 µM and a binding affinity in the low micromolar range. Molecular docking analysis revealed that, in its most stable state, pantinin-1 likely binds to the substrate binding site. The inhibition mode analysis confirmed this result by showing that pantinin-1 acts as a competitive inhibitor. In addition, the cytotoxicity assay demonstrated that pantinin-1 has no toxic effect on Vero cells at concentrations up to 20 µM. However, the potential of pantinin-1 still needs to be tested in a virus cell culture model. Overall, we demonstrated that pantinin-1 could be a promising candidate for the development of antiviral therapy.

## Declaration of generative AI and AI-assisted technologies in the writing process

ChatGPT was used to check grammar and spelling in order to improve readability. After using this tool, the authors reviewed and edited the content and take full responsibility for the content of the publication. Part of the figures were created by BioRender program.

## CRediT authorship contribution statement

**Mohammadamin Mastalipour:** Conceptualization, Methodology, Investigating, Formal analysis, Data curation, Writing – original draft, Writing – review & editing. **Mônika Aparecida Coronado:** Formal analysis, Writing – original draft, Writing – review & editing. **Jorge Enrique Hernández González:** Methodology, Formal analysis, Data curation, Writing – original draft, Writing – review & editing. **Dieter Willbold:** Resources, Writing – original draft, Writing – review & editing. **Raphael Josef Eberle:** Conceptualization, Formal analysis, Supervision, Writing – original draft, Writing – review & editing.

## Acknowledgements

We would like to express our sincere gratitude to Jürgen Manchot Stiftung for providing financial assistance and resources that were critical to the success of this project.

