## Supplement for "Inhibition of Chikungunya virus nsP2 protease *in vitro* by pantinin-1 isolated from scorpion venom"

**Table of content**

Supplementary figure S1. **HPLC chromatograms and mass spectrometry data of pantinin-1**

Supplementary figure S2. Representative BLI sensorgram showing the real-time binding interaction between pantinin-1 and immobilized CHIKV **nsP2^pro^**

Supplementary figure S3. Representative BLI sensorgram showing the control experiment for nonspecific binding of pantinin-1

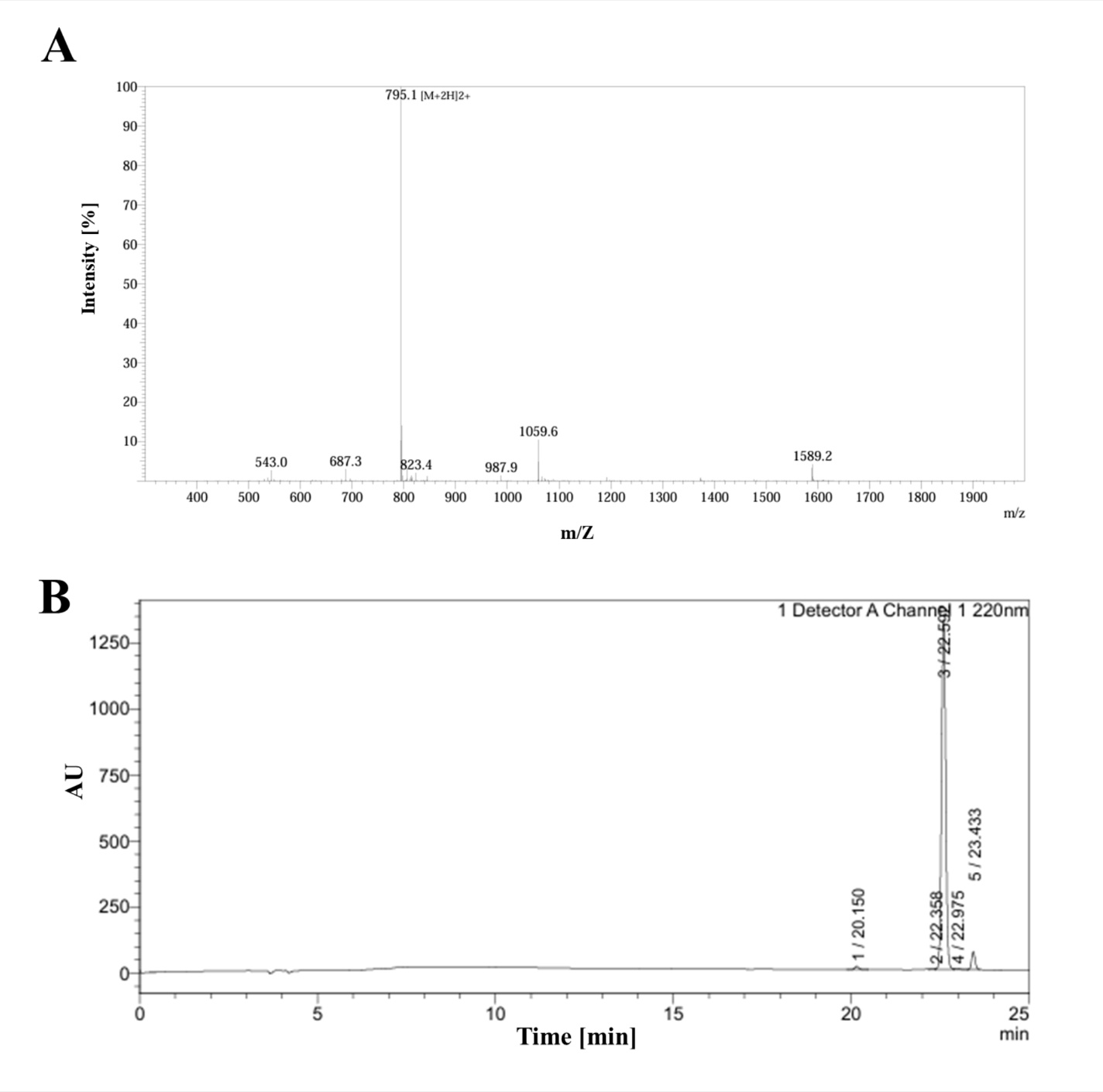

**Fig. S1.** **HPLC chromatograms and mass spectrometry data of pantinin-1.** **(A)** Mass spectrometry data provided by GenScript Biotech (Netherlands) confirming the molecular weight and purity of the synthesized Pantinin-1 peptide. **(B)** HPLC chromatogram showing the retention time and purity profile of pantinin-1.

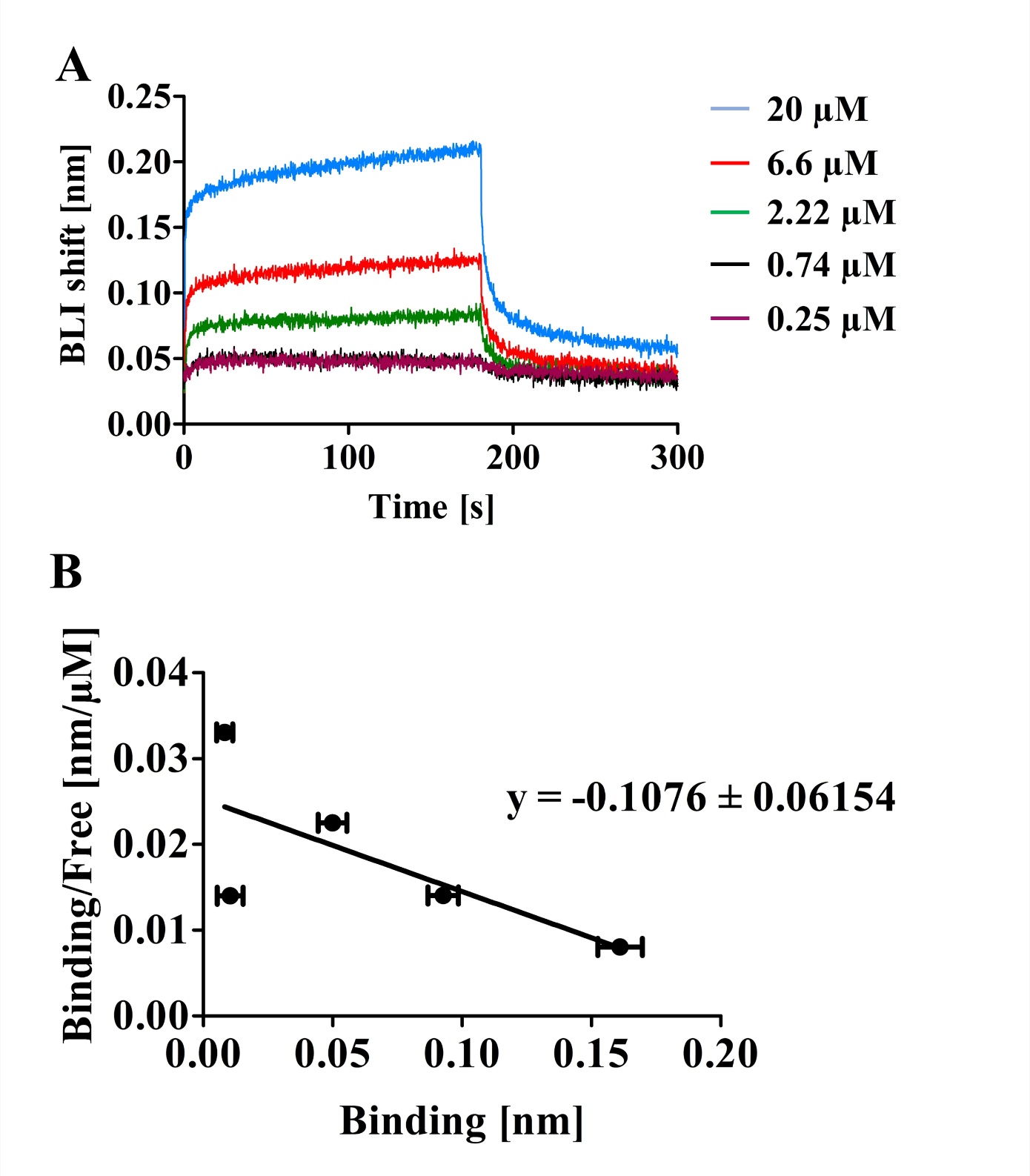

**Fig. S2.** **Fig. S2.** **Representative BLI sensorgram and Scatchard plot analysis of pantinin-1 binding to immobilized CHIKV nsP2^pro^. (A)** Measurement performed using Octet AR2G biosensors. Pantinin-1 was tested at six concentrations (0–20 µM). The sensorgram includes a 180-second association phase followed by a 600-second dissociation phase at room temperature.**(B)** Scatchard plot of pantinin-1 binding to CHIKV nsP2^pro^ based on BLI response data. Binding values at five concentrations were extracted from sensorgrams and plotted as Bound/Free versus Bound. The dissociation constant (K_D_) was calculated from the slope of the linear regression using the equation (K_D_) = –1/slope (Kessler et al., 2008), yielding a (K_D_) of 9.29 µM.

Kessler, M., Suzuki, E., Montgomery, K., & Arai, A. C. (2008). Physiological significance of high- and low-affinity agonist binding to neuronal and recombinant AMPA receptors. *Neurochem Int*, 52(8), 1383-1393. https://doi.org/10.1016/j.neuint.2008.02.00

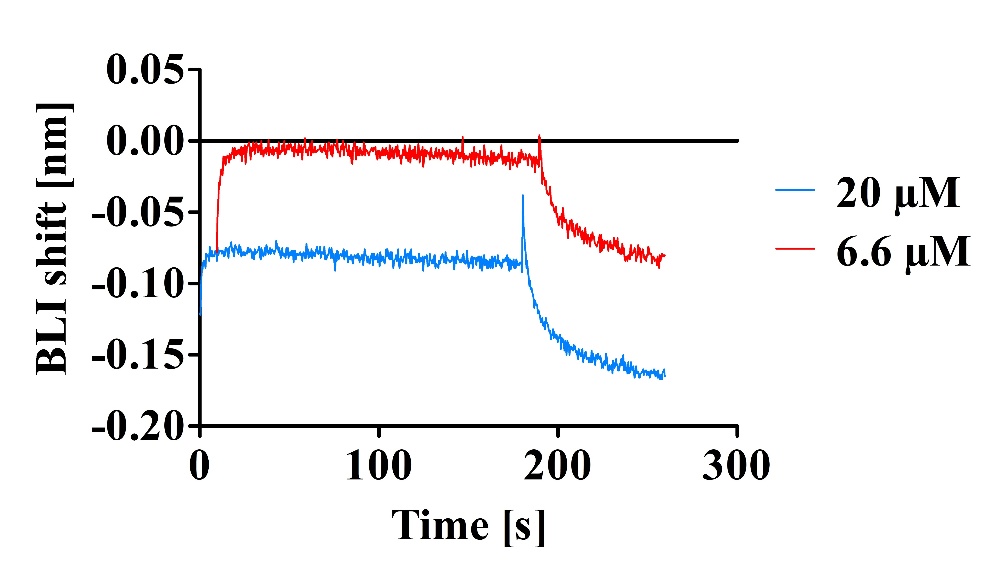

**Fig. S3.** **Representative BLI sensorgram showing the control experiment for nonspecific binding of pantinin-1.** Measurements were performed using Octet AR2G biosensors without immobilized CHIKV nsP2^pro^ to assess non-specific interaction with the sensor surface. Pantinin-1 was tested at two concentrations (20 µM and 6.66 µM). Each sensorgram includes a 180-second association phase and a 600-second dissociation phase at room temperature. No measurable binding response was detected at either concentration, indicating that pantinin-1 does not interact with the biosensor surface in the absence of the target protein.
